# Resolving Cellular Morphology in the Human Brain with Multiparametric Diffusion MR Spectroscopy

**DOI:** 10.1101/2025.10.12.681859

**Authors:** André Döring, Frank Rösler, Kadir Şimşek, Maryam Afzali, Roland Kreis, Karl Landheer, Derek K Jones, Julien Valette, Marco Palombo

## Abstract

Diffusion-weighted magnetic resonance spectroscopy (dMRS) noninvasively probes the diffusion of mostly intracellular metabolites and can therefore report on brain microstructure with a cell-type specificity that water-based diffusion MRI cannot achieve. However, the morphological information accessible to conventional dMRS is limited: estimating both cell-body (soma) and neurite dimensions from a single diffusion-encoding scheme is an ill-posed problem, and neither ultra-high b-value nor diffusion-time–dependent measurements alone distinguish soma size from neurite radius. Here we introduce multiparametric dMRS in the human brain, combining diffusion-time– dependent encoding (apparent diffusion coefficient, ADC, and diffusion kurtosis, K, sampled over diffusion times of 6 ms to 250 ms and b-values up to 8 ms µm^−2^) with double-diffusion-encoded spectroscopy (DDES). Using Monte-Carlo analysis, we show that fitting a two-compartment (soma + neurite) tissue model to diffusion-time data alone is degenerate, admitting two near-indistinguishable solutions. Adding the orthogonal angular information from DDES breaks this degeneracy: jointly fitting both experiments converges to a single, biophysically plausible solution irrespective of initialization, yielding cell-type– specific estimates of intrinsic diffusivity, soma radius, neurite radius, and neurite signal fraction for neuronal (NAA, glutamate) and glial (choline, myo-inositol) metabolites. Metabolite microscopic anisotropy approaches unity, consistent with predominantly intra-neurite diffusion, while simultaneously acquired water data reveal short-range structural disorder and intercompartmental exchange (≈15 ms). Multiparametric dMRS thus extends the standard model of metabolite diffusion by a soma compartment and offers a route toward in vivo, cell-type– specific morphometry of neurons and glia in humans—a foundation for biomarkers in conditions where soma and neurite morphology are altered.

## Introduction

Diffusion-weighted magnetic resonance spectroscopy (dMRS) offers a unique way to noninvasively probe the diffusion of small metabolites in the human brain (1–5). Most brain metabolites are confined to intracellular compartments and preferentially reside in specific cell types; for example, N-acetylaspartate (NAA) and glutamate (Glu) are enriched in neurons, whereas myo-inositol (mI) and choline compounds are abundant in glia(4, 6). By measuring how these metabolites diffuse within their cellular environments, dMRS can provide cell-type–specific insights into microstructure that water-based diffusion MRI, which averages over all cellular and extracellular compartments, cannot resolve.

Despite advances in forward modeling that predict diffusion-MR signals for realistic cellular morphologies, recovering microstructural parameters from measured signals remains difficult. Two factors limit current methods. First, simulating diffusion within realistic cell architectures is computationally expensive, which restricts the use of classical inverse-fitting routines. Second, conventional dMRS experiments sample only a narrow range of diffusion weightings and times, so many models remain underdetermined. Consequently, most human dMRS studies are performed at relatively long diffusion times and high b-values, where sensitive but poorly specific metrics such as the apparent diffusion coefficient (ADC) and diffusion kurtosis (K) can be estimated robustly (7).

The time scale of diffusion encoding determines which structural features are emphasized. Pulsed-gradient (PG) sequences, used at long diffusion times (TD ≥ 50 ms), primarily report on diffusion along tortuous cellular processes (8–10). In contrast, oscillating gradients (OG) enable shorter diffusion times (≤10 ms) and increase sensitivity to smaller structures such as somata, subcellular organelles, and fiber diameters (11, 12). Practical constraints on human scanners— peripheral nerve stimulation (PNS), gradient slew limits, and acoustic resonances—cap the achievable b-value at very short diffusion times, so in that regime only the ADC can be measured reliably. Nevertheless, sampling the ADC across a broad range of times, from milliseconds to hundreds of milliseconds, provides access to microstructural information across multiple length scales.

Diffusion-encoding MRS sequences can be categorized by their targeted diffusion times, with stimulated-echo sequences (e.g., STEAM) applied with PG at long diffusion times and spin-echo sequences (e.g., semiLASER, PRESS) at short (with OG) to intermediate (with PG) diffusion times (5, 13– 15). Figure 1a gives a representative overview of the sequences and diffusion-encoding strategies for the corresponding diffusion-time intervals.

**Fig. 1:**
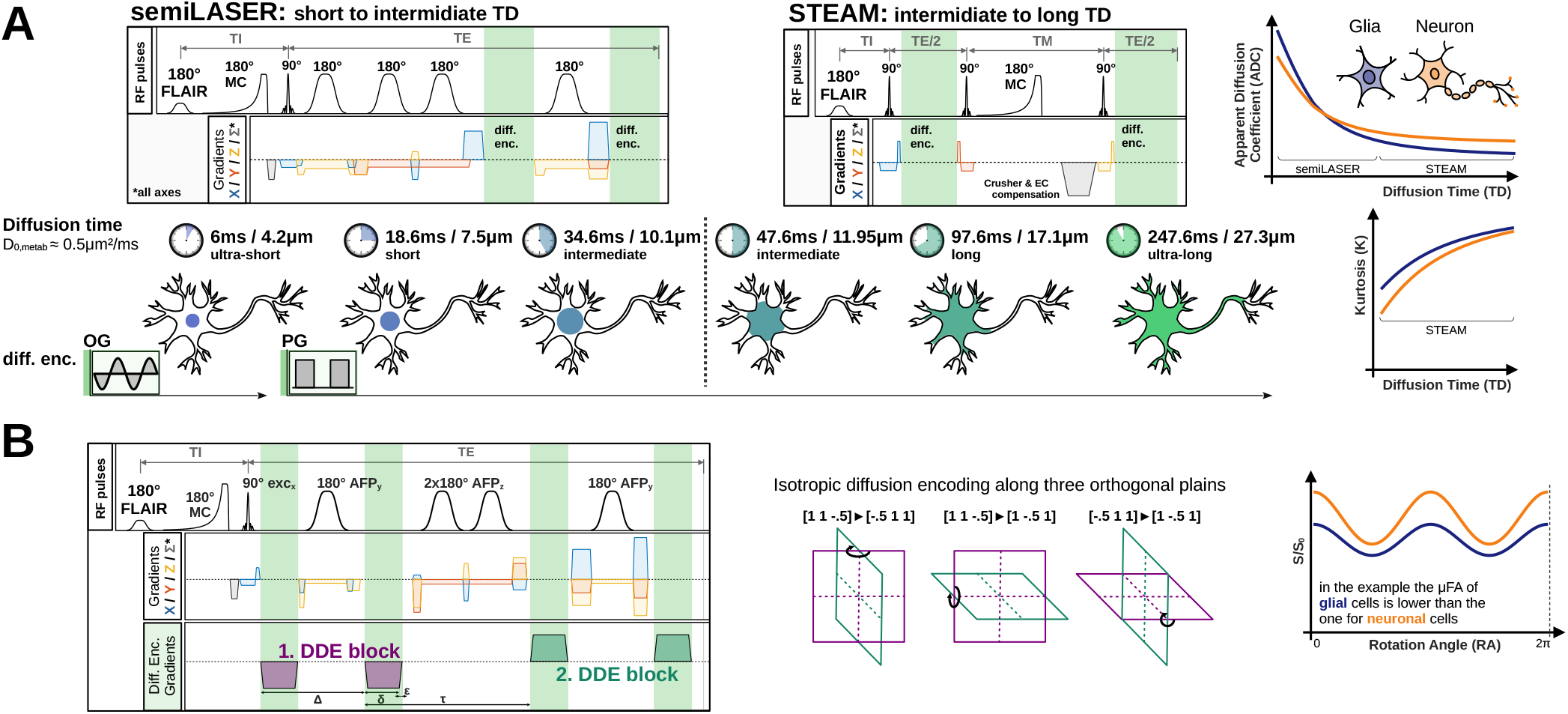
Acquisition strategies. (A) STEAM and semiLASER sequences were used to measure the diffusion-time (TD) dependence of the apparent diffusion coefficient (ADC) and diffusion kurtosis (K). In the short-TD range (6, 21, 37 ms), semiLASER was used up to a b-value of 3 ms µm^−2^ with oscillating gradients (OG) at the shortest TD and pulsed gradients (PG) otherwise. In the long-TD range (50, 100, 250 ms), STEAM was used up to a b-value of 8 ms µm^−2^ with PG encoding. (B) The semiLASER sequence was extended with double-diffusion-encoding (DDE) gradient pairs applied in three orthogonal planes to achieve powder-averaged diffusion encoding. Both sequences are equipped with metabolite cycling to measure water and metabolite diffusion simultaneously, and with fluid-attenuated inversion recovery (FLAIR) to suppress cerebrospinal fluid (CSF).

To further extend the sampled parameter space, complementary information can be obtained by applying two successive diffusion-encoding periods of variable relative orientation. This technique, double-diffusion encoding (DDE), enables two-dimensional encoding of diffusion and can distinguish microscopic compartments such as cylinders and spheres even in macroscopically disordered tissue (16, 17). Spectroscopic double-diffusion-encoded experiments (DDES) combine DDE with spectroscopic localization, here implemented in a semiLASER sequence (Figure 1b). Because DDE probes the angular dependence of diffusion, it provides additional constraints on cell geometry and may help to discriminate neurite from soma contributions, independent of diffusion time.

The standard model of metabolite diffusion currently consists of randomly oriented neurites alone, modeled as thin cylinders or sticks, and even this simplified model requires ultra-high b-values (> 10 ms µm^−2^) to be estimated reliably from a single encoding scheme (5, 18, 19). Extending this model with a second, soma-like compartment is conceptually analogous to the soma-and-neurite density imaging (SANDI) framework developed for water diffusion (20), but transferring such a two-compartment description to metabolites—and resolving its parameters in vivo—has not previously been achieved in humans. So far, multiparametric dMRS combining diffusion-time and angular encoding has been demonstrated only in animals (21).

In this study we implement a multiparametric dMRS protocol in humans that combines diffusion-time–dependent measurements with DDES. We acquire ADCs over diffusion times of 6 ms to 250 ms, extend the b-value range up to 8 ms µm^−2^ to estimate kurtosis at diffusion times ≥50 ms, and acquire DDES in an eight-step rotation cycle at a b-value of 5.2 ms µm^−2^. The resulting dataset spans short to long diffusion times and includes angular dependence, enabling us to fit two-compartment tissue models that separately estimate soma radius (R_s_), neurite radius (R_c_), neurite signal fraction (f_c_), and intrinsic diffusivity (D_0_). We further model the DDES data with a diffusion-tensor representation to derive microscopic fractional anisotropy (*μ*FA), axial (D_∥_ ) and radial (D_⊥_) diffusivities, and mean diffusivity (MD).

Our objectives are threefold: (i) to demonstrate that the diffusion-time dependence of metabolite ADC and kurtosis can be measured in humans; (ii) to evaluate whether DDES provides complementary information on cellular geometry; and (iii) to test whether jointly fitting diffusion-time– dependent and DDES data improves the stability and physical plausibility of microstructural estimates. To this end, we use a ultra-strong-gradient Connectom scanner and translate the DDES protocol to conventional clinical 3 T systems. By integrating diffusion-time and angular information, we aim to overcome the ambiguities inherent in single-encoding models and to provide a framework for cell-type–specific characterization of neuronal and glial morphology in vivo.

## Materials and Methods

### Participants, ethics, and study design

This study combined two complementary experiments acquired in two independent cohorts, reflecting the differing hardware requirements of the two encoding schemes. Diffusion-time experiments, which require ultra-strong gradients to reach short diffusion times, were performed on a Connectom scanner; DDES experiments, which are feasible on clinical hardware, were performed on conventional 3 T systems. All participants were screened for MR contraindications and gave written informed consent. The diffusion-time cohort comprised ten healthy adults ((32.8 ± 3.4); seven female), recruited with approval from the Cardiff University School of Psychology Research Ethics Committee. The DDES cohort comprised four healthy adults ((56.7 ± 14.4) ; all male), recruited in agreement with the local ethics review boards at the participating sites.

### Anatomical imaging and shimming

Voxel positioning and tissue segmentation used 1 mm^3^ isotropic MPRAGE images (TE /TI /TR = 2 /857 /2300 ms, flip angle 9^°^). First- and second-order shimming used the vendor’s automated double-acquisition gradient-echo routine (Siemens Healthineers, Erlangen, Germany).

### Diffusion time experiments

Measurements were conducted on a 3 T Connectom-A MR scanner (Siemens Healthineers) with a 32-channel receive head coil. Metabolite diffusion was measured at (i) short-to-intermediate TDs (6 ms, 21 ms and 37 ms) using semiLASER (b_max_ = 3 ms µm^−2^, G_max_ = 254 mT m^−1^) (14), and (ii) intermediate-to-long TDs (50 ms, 100 ms and 250 ms) with STEAM (b_max_ = 8 ms µm^−2^, G_max_ = 151 mT m^−1^) (22); detailed settings are in Supplementary Table S2. Diffusion gradients had a bipolar sinusoidal shape at the shortest TD and a trapezoidal shape otherwise (11, 14). For STEAM, six outer-volume-suppression (OVS) bands surrounded the volume of interest (VOI) to improve localization and suppress lipid contamination and spurious echoes. The diffusion-encoding direction was set along the *x* /*y* diagonal, with identical gradient amplitudes to reach sufficiently high b-values while avoiding the *z* axis to minimize table vibration. For *b* ≤ 3 ms µm^−2^, 32 transients were acquired, and 64 otherwise. FLAIR was used to suppress CSF, and metabolite cycling to measure water and metabolite diffusion simultaneously. Cross terms were compensated by inverting the diffusion-gradient polarity in successive transients. For STEAM, a macromolecular baseline (MMBG) was acquired at all TDs by metabolite nulling (inversion time 765 ms) with moderate diffusion weighting (5 ms µm^−2^); the MMBG was parametrized and smoothed with a model of 80 Voigt lines (23), and the residual Cr_CH2_ moiety was removed by modeling and subtracting a single Voigt line (T_2_ = 125 ms) at 4.0 ppm.

The VOI ((10.0 ± 1.5) mL) was placed midline in the posterior cingulate cortex (PCC), maximizing gray-matter fraction (mean (70.9 ± 1.7) %) while avoiding subcutaneous fat and CSF. The number of repeated acquisitions per TD (in ms) was: {6: 6}, {21: 3}, {37: 7}, {50: 8}, {100: 7}, {250: 8}.

### Double-diffusion encoded experiments

Measurements were conducted at two sites on 3 T Siemens PRISMA MR scanners with 32-channel receive head coils. Non-water-suppressed diffusion-weighted spectra were acquired with the same semiLASER sequence used for the diffusion-time experiments (14). The crushing scheme was optimized to reduce spurious diffusion weighting by splitting crusher gradients to null the zeroth-order moment between RF pulses (Figure 1b). Diffusion weighting was applied at *b* = 5.2 ms µm^−2^ with DDE encoding along three orthogonal planes ([1.0, 1.0, −0.5], [−0.5, 1.0, 1.0], [1.0, − 0.5, 1.0] ) using an eight-step rotation cycle (0, 45, …, 315^°^). Interleaved within the rotation cycle, 3 × 16 transients were acquired at *b* = 0. For each rotation angle, positive- and negative-polarity data were acquired with 24 transients each (144 per angle). Total acquisition time was 70 min per participant (Supplementary Table S2). The VOI ((16.1 ± 2.4) mL) was placed midline in the PCC (gray-matter fraction (63.0 ± 5.4) %).

### Data processing

Single transients were stored in raw twix format without coil-channel averaging or preprocessing. A custom MATLAB pipeline (MathWorks) performed coil-channel combination, phase-offset, frequency-drift, and eddy-current correction, and motion compensation based on the inherent water reference (13, 24).

Metabolite basis sets were simulated in MARSS, accounting for sequence type (spin/stimulated echo), timings (TE and TM), and RF-pulse profiles (25). The fitting model comprised 18 metabolites: aspartate (Asp), *γ*-aminobutyric acid (GABA), glucose (Glc), glutamine (Gln), Glu, glycine (Gly), glutathione (GSH), lactate (Lac), mI, NAA, N-acetylaspartylglutamate (NAAG), phosphorylethanolamine (PE), scyllo-inositol (sI), taurine (Tau), total choline [tCho = glycerophosphocholine (GPC) + phosphocholine (PCho)], and total creatine [tCr = creatine (Cr) + phosphocreatine (PCr)]. To account for differing T_2_ values of metabolite moieties, relative Lorentzian broadenings were applied: 0.15 Hz to NAA_asp_, 0.50 Hz to NAAG_asp_ and NAAG_glu_, 0.70 Hz to the Cr_CH2_ and PCr_CH2_ singlets, 0.20 Hz to the PCho and GPC singlets, and 0.20 Hz to the GSH singlet (26). For STEAM, the parametrized MMBG patterns were added to the fitting model at each TM (23).

Spectral fitting was performed by linear-combination modeling in FiTAID using Levenberg–Marquardt *χ*^2^-minimization in the frequency domain over 0.0–4.2 ppm (27, 28). Metabolites within the groups [Asp, NAAG, PE, tCr], [Glc, mI, sI], [Gln, Glu], and [Gly, GSH] were constrained to equal Lorentzian broadening, while frequency offset, zeroth-order phase, and Gaussian broadening were shared across all metabolites. Effective b-values were derived from the sequence chronograms, accounting for crusher and slice-selection gradients.

### Sequence validation

Both sequences were validated in two room-temperature phantom experiments ((20.5 ± 0.7)°C); the full analysis is in Supplementary Figure S1. First, a National Institute of Standards and Technology (NIST) phantom containing a 40 % mass fraction of polyvinylpyrrolidone (PVP), used to slow diffusion, served to exclude bias related to diffusion-encoding strategy (OG vs. PG) and diffusion time; all settings matched the in-vivo condition, and free-water diffusion coefficients were estimated by mono-exponential fitting. Second, a “BRAINO” phantom (GE HealthCare) was used to measure metabolite diffusion and exclude bias from differences in basis sets and from J-evolution when TE, TM, echo type, or diffusion-encoding scheme vary. Because of fast metabolite diffusion, the b-value range was reduced to eight linearly spaced values from 0.1 to 1.5 ms µm^−2^. Diffusion-weighted spectra of mI (5.0 mM), NAA (12.5 mM), Glu (12.5 mM), Cr (10.0 mM), Lac (5.0 mM), and Cho (3.0 mM) were fitted in FiTAID, and pattern amplitudes informed a mono-exponential free-diffusion model and a diffusion-tensor representation in MATLAB to estimate individual metabolite diffusivities. Validation results are summarized in Section “Spectral quality” and Supplementary Figure S1.

### Two-compartment tissue model

Tissue models were implemented in MATLAB using analytical solutions for PG and OG encoding (29–31) and the matrix formalism (MISST toolbox) for DDE encoding (32–36). The model comprised two non-exchanging compartments: randomly oriented (powder-averaged) cylinders representing neurites—hereafter “astro-cylinders”—and spheres representing somata. Its parameters are D_0_ (intrinsic free diffusivity), R_c_ (cylinder radius, reflecting effective neurite radius), R_s_ (sphere radius, reflecting effective soma size), and f_c_ (cylinder signal fraction, with f_s_ = 1 − f_c_ the sphere fraction).

### Diffusion-time representations and model fitting

Metabolite areas and their Cramér–Rao lower bounds (CRLBs)—used as inverse weights—informed two diffusion representations: (i) mono-exponential fitting of ADC_E_ (TD) at low b-values (≤ 3 ms µm^−2^) for both semiLASER and STEAM, and (ii) a kurtosis representation, ADC_K_ (TD) and K (TD), for the STEAM acquisitions over the full b-value range. An outlier-detection step removed spectra potentially affected by motion: either all b-values were retained or one b-value was iteratively removed, selecting the fit with the lowest corrected Akaike information criterion (AICc). For microstructural modeling, only the diffusion-time dependence of ADC_E_ (TD) and K (TD) was used.

For a given configuration (D_0_, R_s_, R_c_, f_c_ ), the signal was simulated for each TD and up to b_max_ = 8 ms µm^−2^ using the b-values applied in the STEAM experiments. The model-specific mono-exponential ADC_E_ and kurtosis K were obtained by least-squares fitting up to 3 ms µm^−2^ and 8 ms µm^−2^, respectively, yielding the model functions ADC_E_(D_0_, R_s_, R_c_, f_c_, TD, b) and K(D_0_, R_s_, R_c_, f_c_, TD, b). These were fitted simultaneously to the measured ADC_E_(TD) and K (TD) using MATLAB’s trust-region-reflective algorithm, with 0.40 ≤ f_c_ ≤ 0.90. Fitting uncertainties were estimated by residual bootstrapping with 250 noise realizations.

### Monte-Carlo stability analysis

To assess fitting stability and identifiability, we performed Monte-Carlo (MC) simulations with 10 000 uniformly distributed starting parameters in the ranges 0.00 ≤ D_0_ ≤ 0.50 µm^2^ ms^−1^, 0 ≤ R_c_ ≤ 15 µm, 0 ≤ R_s_ ≤ 40 µm, and 0.40 ≤ f_c_ ≤ 0.90. The median of the lowest-RMSE 10% quantile of fits constrained to R_c_ < R_s_ provided physically plausible starting values for refitting the diffusion-time dependence (Supplementary Figure S2). We also tested a joint formulation across metabolites inhabiting the same compartment, with metabolite-specific D_0,i_ but shared R_s_, R_c_, and f_c_, giving ADC_E_(D_0,i_, R_s_, R_c_, f_c_, TD, b) and K(D_0,i_, R_s_, R_c_, f_c_, TD, b), with *i* the metabolite index.

### DDES modeling

The DDES experiment was modeled in two ways: (i) the two-compartment tissue model (simulated using MISST), accounting for all rotation-angle configurations, giving DDE (D_0_, R_s_, R_c_, f_c_, Θ, b); and (ii) a signal representation fitting the powder average of 256 uniformly oriented diffusion tensors, giving DDE (D_∥_, D_⊥_, Θ, b). The estimated axial (D_∥_) and radial (D_⊥_ ) components were used to compute the mean diffusivity (MD = (D_∥_ + 2D_⊥_) /3) and the microscopic fractional anisotropy (*μ*FA) (17).

### Joint multiparametric modeling

A joint model combined ADC_E_ (D_0_, R_s_, R_c_, f_c_, TD, b), K (D_0_, R_s_, R_c_, f_c_, TD, b), and DDE (D_0_, R_s_, R_c_, f_c_, Θ, b) to fit the diffusion-time and DDES data simultaneously with a single set of microstructural parameters. Model complexity and long computation times precluded full MC initialization; instead, model stability was probed by initializing from four sets of starting conditions drawn from the MC results: the optimal R_c_ < R_s_ solution and three further sets far from it (including a non-physical R_c_ > R_s_ solution; Supplementary Table S3).

### Modelling of water diffusion

Water was acquired simultaneously with metabolites by metabolite cycling, using identical settings. For the diffusion-time experiment, a kurtosis representation (ADC_K,H2O_(TD) and K_H2O_(TD)) was fitted to b-values ≤ 3 ms µm^−2^ for both semiLASER and STEAM. We then fitted (i) a power-law model jointly to ADC(TD) and K(TD) to estimate structural disorder, (ii) a Kärger model to K(TD) alone to estimate the exchange time, and (iii) a diffusion-tensor model to the water DDES signal DDE(Θ) to estimate *μ*FA, D_∥_, and D_⊥_.

## Results

### Spectral quality

Figure 2 shows representative diffusion-weighted spectra from the diffusion-time and DDES experiments in a single participant with the VOI placed midline in the PCC. In both cases, spectra are free of eddy-current artifacts and residual water, with overall quality consistent with consensus recommendations (5). Linear-combination modeling left only minor residuals, largest for the major singlets at the lowest b-value.

**Fig. 2:**
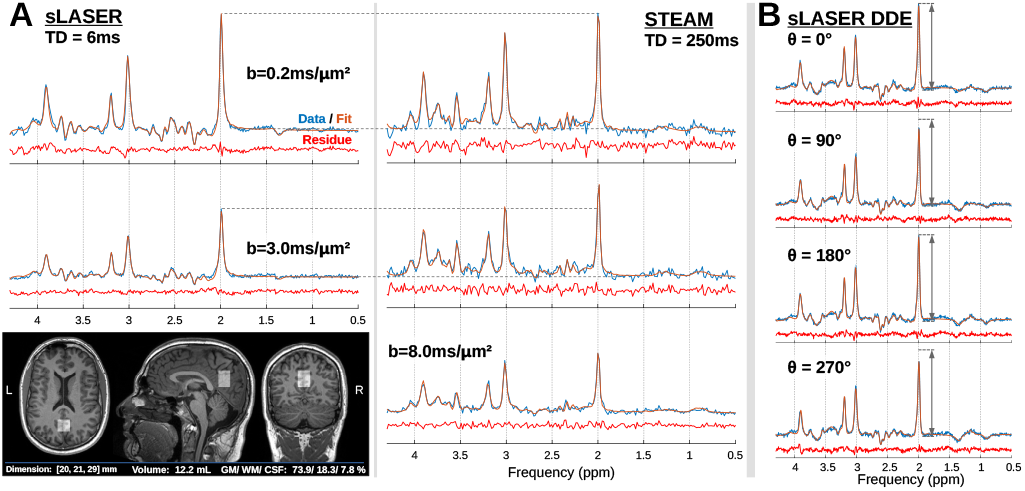
Spectral quality and fitting. (A) Spectral fitting at the shortest TD (6 ms, semiLASER; *b* = 0.2 and 3.0 ms µm^−2^ ) and the longest TD (250 ms, STEAM; *b* = 0.2, 3.0, and 8.0 ms µm^−2^ ); inset shows the voxel position. Spectra at the lowest b-value are scaled to equal NAA peak height, with faster NAA diffusion at the shortest TD evident from stronger attenuation. (B) DDES fitting at four rotation angles; NAA peak height is referenced to Θ = 0°, showing the modulation of NAA intensity. Linear-combination modeling (orange) and residuals (red) indicate good fit quality throughout.

For the diffusion-time experiment (Figure 2a), spectra at the shortest and longest TDs (6 and 250 ms) are shown. The cohort-wide signal-to-noise ratio (SNR), relative to the NAA peak at *b* = 3 ms µm^−2^, was 36.8 ± 9.0 for semiLASER and 45.8 ± 6.8 for STEAM, above the minimum consensus recommendation for dMRS (5). The NAA linewidth increased slightly toward higher b-values, with means of (4.7 ± 0.6) Hz (semiLASER) and (4.5 ± 0.5) Hz (STEAM), averaged over all b-values, TDs, and participants. Spectra at 6 and 250 ms are scaled to equal NAA peak height at the lowest b-value; the faster NAA diffusion at TD = 6 ms is directly evident from the stronger attenuation at *b* = 3 ms µm^−2^ (other peaks are not directly comparable because of metabolite-specific T_2_/T_1_ and TE/TM differences).

For DDES (Figure 2b), the oscillatory diffusion signal is directly visible across the four shown rotation angles (0–270°), with the largest signal for the parallel/antiparallel configuration (0/180°) and the smallest for the perpendicular configuration (90/270°). A high mean SNR at *b* = 5.2 ms µm^−2^ (56.4 ± 16.6, averaged over all rotation angles) enabled stable fitting of the subtle sinusoidal modulation; the NAA linewidth ((4.7 ± 0.7) Hz) matched the diffusion-time experiments.

Moreover, phantom validation confirms that estimates were free of sequence-related bias (Supplementary Figure S1). In the NIST phantom, signal attenuation was mono-exponential and the estimated free-water diffusivity was consistent across sequence type, echo time, and encoding scheme (mean (0.56 ± 0.02) µm^2^ ms^−1^). In the BRAINO phantom, free metabolite diffusivities were likewise independent of echo, mixing, and diffusion time—NAA 0.57 ± 0.04, tCr 0.74 ± 0.04, Cho 0.84 ± 0.04, Lac 0.81 ± 0.04, Glu 0.64 ± 0.06, and mI 0.58 ± 0.15 µm^2^ ms^−1^—and followed an inverse power-law relationship with molecular weight (exponent *α* = 0.58 ± 0.09, *p* < 0.01), in line with prior work (37). DDES validation in the BRAINO phantom yielded *μ*FA≈0 for all metabolites and D_∥_ ≈ D_⊥_ ≈ MD independent of encoding direction, with mean diffusivities matching the diffusion-time experiments.

### Diffusion-time dependence of ADC and kurtosis

Figure 3 compares our estimated ADC_E_ (TD) and K (TD) to values reported in humans and animals. Mono-exponentially fitted metabolite ADCs were highest at the shortest TD (6 ms), ranging from 0.15 to 0.29 µm^2^ ms^−1^, and plateaued for TD ≥ 50 ms between 0.09 and 0.11 µm^2^ ms^−1^ at the longest TD (250 ms). With the exception of Glu, kurtosis increased from 1.0–1.4 at TD = 50 ms to 1.6–2.0 at TD = 250 ms (metabolite-specific values in Supplementary Table S4). The decrease in Glu kurtosis at the longest TD is likely related to its low SNR at high b-values and long TD.

**Fig. 3:**
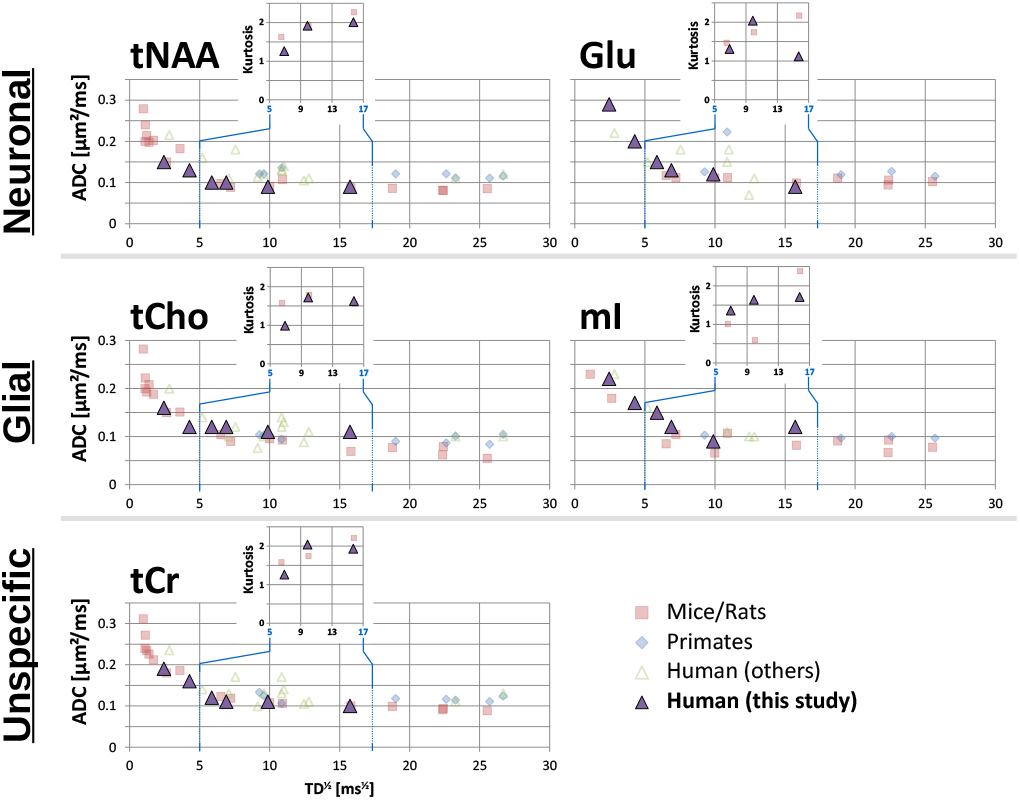
Literature comparison. Diffusion-time dependence of ADC_E_ (TD) and K(TD) for selected brain metabolites, compared with prior studies in rodents (□), nonhuman primates (⋄), and humans (△). Data sources: Najac et al. (9), Ligneul and Valette (11), Döring et al. (13), Döring and Kreis (14), Mougel et al. (21), Pfeuffer et al. (38), Ellegood et al. (39), Valette et al. (40), Kan et al. (41), Marchadour et al. (42), Najac et al. (43), Ercan et al. (44), Palombo et al. (45), Deelchand et al. (46), Ingo et al. (47).

### Diffusion-time–only modeling is degenerate

Monte-Carlo simulations of the two-compartment model revealed an ill-posed problem with an unstable, ambiguous solution space (Supplementary Figure S2). RMSE values spanned several orders of magnitude, and two near-indistinguishable solution branches emerged, with almost identical fitting curves over the measured TD range: (i) a non-physical branch with R_c_ > R_s_ (larger neurite than soma radius), reached at marginally lower RMSE, and (ii) a physically plausible branch with R_c_ < R_s_. The two branches differed mainly in the back-extrapolation of K (TD) toward short, unmeasured diffusion times. This degeneracy was consistent across all metabolites.

Obtaining a physically meaningful solution from diffusion-time data alone therefore required constraining the solution space to R_c_ < R_s_ and careful, physically informed initialization from the MC results (Supplementary Figure S2). The resulting intrinsic diffusivities and microstructural parameters are reported in Table 1, with bootstrap confidence intervals. The intrinsic diffusivities D_0_ varied most, alongside R_s_, whereas R_c_ and f_c_ were comparatively consistent across metabolites. D_0_ was highest for NAA, Glu, mI, and tCr, and lowest for tNAA and tCho. Soma radii aligned for NAA, tNAA, mI, and tCr (mean 7.0 µm) but reached large values for Glu (20.4 µm) and tCho (38.3 µm). Neurite radii ranged from 1.2 to 2.5 µm, and neurite fractions f_c_ were consistent (0.82–0.89), reaching the upper boundary for Glu and the joint fits. Although the joint (multi-metabolite) formulation lowered the AICc, the estimated uncertainties were unchanged and the large soma radii of Glu and tCho propagated into the joint fits.

**Table 1:** Diffusion-time experiment. Cellular microstructural parameters (R_s_, R_c_, f_c_) and intrinsic diffusivities (D_0_ ) constrained to R_c_ < R_s_. Yellow marks fits that converged to the boundary condition f_c_ = 0.90.

|  | NAA | tNAA | Glu | NAA+tNAA* | NAA+tNAA*+Glu | tCho | mI | tCho+mI | tCr |
| --- | --- | --- | --- | --- | --- | --- | --- | --- | --- |
| $D_0$<br>[$\mu\text{m}^2/\text{ms}$] | 0.45±0.03 | 0.33±0.02 | 0.42±0.06 | 0.49±0.03<br>0.32±0.02 | 0.35±0.04<sup>NAA</sup><br>0.24±0.02<sup>tNAA</sup><br>0.44±0.05<sup>Glu</sup> | 0.24±0.02 | 0.40±0.03 | 0.25±0.02<sup>tCho</sup><br>0.33±0.02<sup>mI</sup> | 0.44±0.04 |
| $R_s$ [$\mu\text{m}$] | 7.4±0.7 | 6.2±0.7 | 20.4±8.5 | 6.6±0.5 | 33.4±1.2 | 38.3±1.5 | 8.5±1.7 | 34.9±1.8 | 5.9±0.9 |
| $R_c$ [$\mu\text{m}$] | 1.5±0.2 | 1.2±0.2 | 2.5±0.5 | 1.3±0.1 | 2.1±0.1 | 2.4±0.3 | 2.0±0.2 | 2.4±0.1 | 1.3±0.2 |
| $f_c$ | 0.82±0.04 | 0.82±0.02 | 0.90±0.07 | 0.83±0.02 | 0.90±0.03 | 0.86±0.05 | 0.89±0.04 | 0.90±0.03 | 0.82±0.04 |
| AICc | -4.6 | -3.9 | -2.4 | -9.1 | -11.8 | -3.7 | -4.1 | -8.6 | -3.5 |

\*: for joint fitting of metabolites in the neuronal compartment, tNAA was used instead of NAA alone because of low NAA SNR and distinct $D_0$ differences between NAA and tNAA..

### Double-diffusion encoded spectroscopy

Figure 4 shows DDES fitting for a single participant. The oscillatory signal DDE (Θ) was clearly resolved for NAA, tNAA, tCho, and tCr, and less pronounced for Glu and mI, which also had the largest CRLBs. The two-compartment tissue model and the tensor representation gave consistent results, with D_∥_ close to the intrinsic diffusivity D_0_ for all metabolites.

**Fig. 4:**
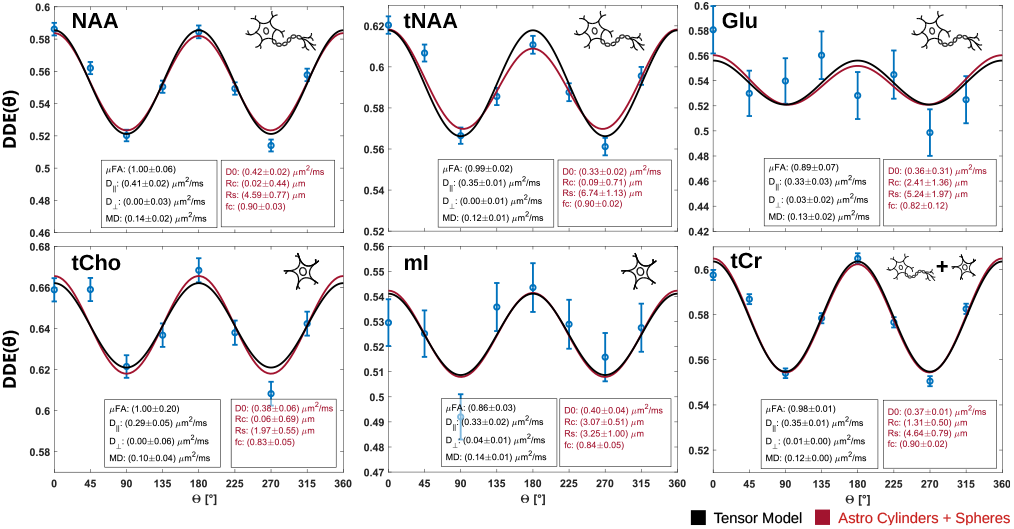
DDES fitting. Oscillatory diffusion signal for a single participant, fitted with two models (tissue model: astro-cylinders + spheres; tensor model). Error bars are derived from CRLBs; microstructural metrics (insets) are reported with bootstrap standard deviations.

Cohort results are summarized in Table 2 (two participants excluded for Glu and three for mI because of poor oscillatory-signal quality). The cohort-wide intrinsic diffusivities D_0_ and axial diffusivities D_∥_ agreed with the diffusion-time estimates constrained to R_c_ < R_s_. Mean diffusivities were lower by a factor of 3–4 (0.10–0.14 µm^2^ ms^−1^), consistent with a compartment partly composed of randomly oriented cylinders; indeed, MD ≈ D_0_ /3, the expected signature of stick-like diffusion. Radial diffusivities D_⊥_ were close to zero for all metabolites, indicating limited sensitivity to small neurite radii—most likely due to coarse-graining at the relatively long DDES diffusion time of 27.3 ms (10). Correspondingly, neurite radii R_c_ tended to zero for most metabolites (except Glu and mI). Soma radii R_s_ (3.3–6.8 µm) were consistent across metabolites and larger in neurons than in glia, and neurite fractions f_c_ aligned across metabolites (0.83–0.90). The microscopic fractional anisotropy *μ*FA was close to 1.0 for all metabolites, indicating a highly anisotropic environment on the ≈10 µm length scale probed here—the expected signature of predominantly intra-neurite (stick-like) diffusion.

**Table 2:**
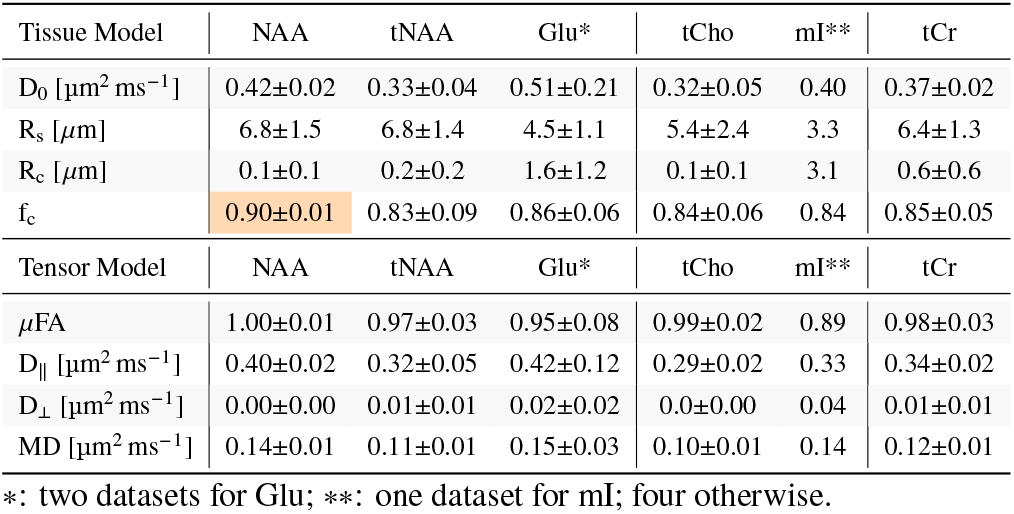
DDES experiments. Cellular microstructural parameters (R_s_, R_c_, f_c_, D_0_ ) from the two-compartment model and diffusion metrics (*μ*FA, D_∥_, D_⊥_, MD) from the tensor model.

### Joint multiparametric modeling resolves the degeneracy

Jointly fitting the diffusion-time and DDES data (Figure 5) resolved the degeneracy seen with diffusion-time data alone. For every metabolite, the fit converged to the same bio-physically plausible solution (R_c_ < R_s_) irrespective of the four very different starting conditions, giving largely overlapping curves for ADC (TD), K (TD), and DDE (Θ) . The standard deviation across the four converged fits was small, reflecting the reproducibility of convergence rather than measurement uncertainty (see caption). Intrinsic diffusivities D_0_ were highest for NAA, Glu, mI, and tCr (0.44–0.48 µm^2^ ms^−1^) and lowest for tNAA and tCho (0.34 and 0.35 µm^2^ ms^−1^). Neurite radii were ≈1 µm, highest for Glu. The mean soma radius of the neuronal markers NAA and tNAA (6.8 µm) was slightly smaller than that of the glial markers tCho and mI (9.0 µm). Notably, the implausibly large soma radius found for tCho in the diffusion-time–only fit stabilized to 9.8 µm in the joint fit, whereas the large Glu soma radius persisted (20.6 µm); tCr had the smallest soma radius (6.0 µm). Neurite fractions were equal for NAA, tNAA, tCho, and tCr, and reached the boundary for Glu and mI.

**Fig. 5:**
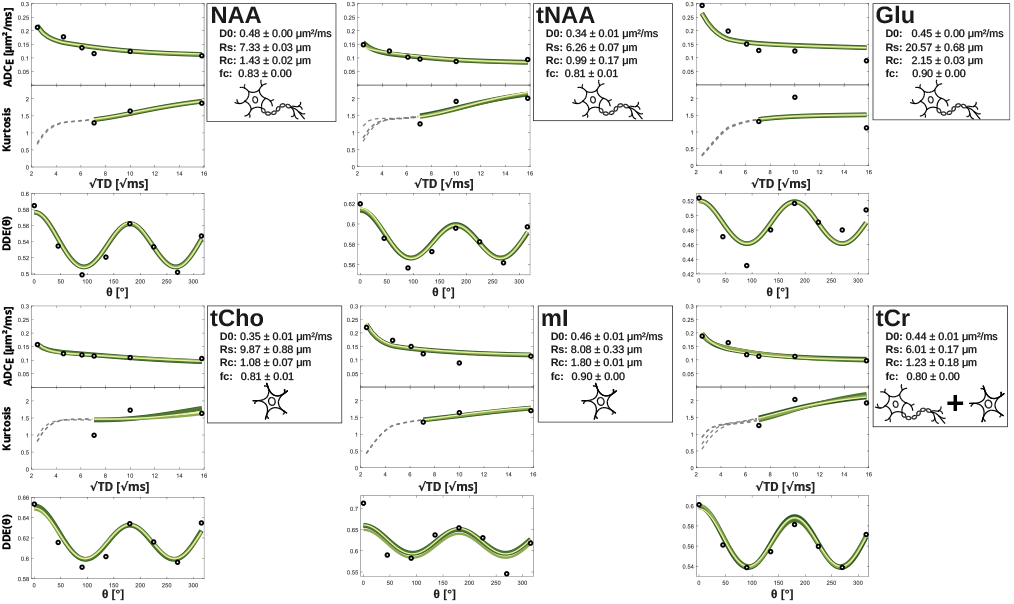
Joint fitting of diffusion-time and DDES data. Cohort-averaged data fitted with the two-compartment model (astro-cylinders + spheres). Four sets of starting conditions were used to probe stability; the resulting ADC_E (_TD), K (TD), and DDE (Θ) curves are color-coded, with dashed lines showing the back-extrapolation of K (TD) toward short diffusion times. Insets give microstructural parameters (D_0_, R_s_, R_c_, f_c_). The reported uncertainties are the standard deviation across the four converged fits (convergence reproducibility), not a measurement uncertainty.

### Water diffusion and exchange

Water, acquired simultaneously by metabolite cycling, provided complementary microstructural information (Figure 6). The diffusion-time dependence of ADC_K_ (TD) did not decay monotonically but remained highest at the shortest TD (ADC_K,6_ = (1.12± 0.08) vs. ADC_K,250_ = (1.00 ± 0.09 µm^2^ ms^−1^). In contrast, kurtosis was lowest at the shortest TD, peaked at TD^∗^ = 21 ms (0.65 ± 0.01), and then declined toward longer TDs—a characteristic signature of exchange, weak at short TD and dominant at long TD (48–51).

**Fig. 6:**
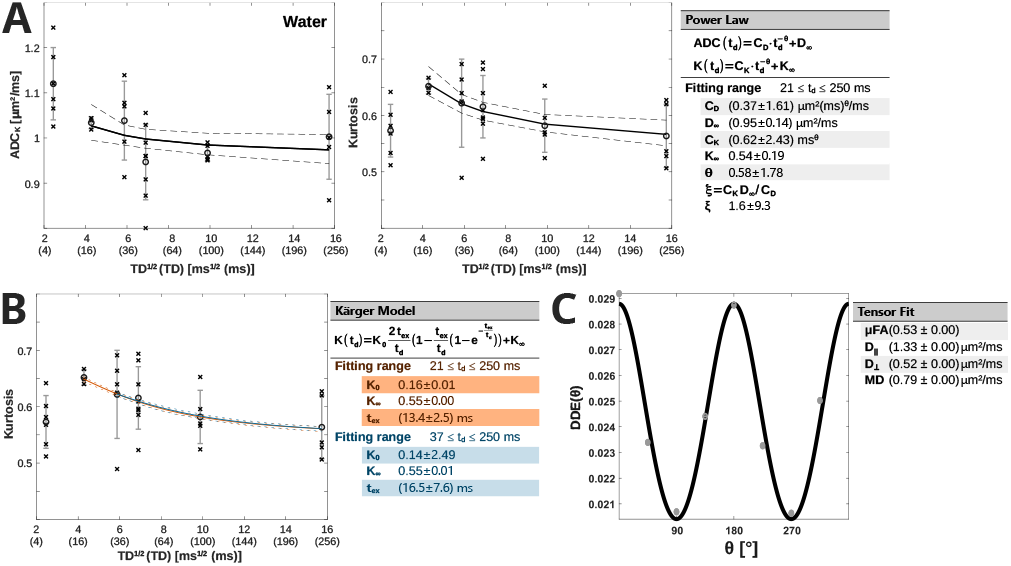
Water diffusion. (A) Joint fitting of ADC(TD) and K(TD) with a power-law model. (B) Fitting of K(TD) with a Kärger exchange model over two restricted TD ranges. (C) Fitting of the water DDES signal DDE(Θ) with a diffusion-tensor model.

A power-law model fitted jointly to ADC (TD) and K (TD) over 21 ≤ TD ≤ 250 ms (the range of model validity, requiring monotonic K (TD) decay) (52, 53) returned dynamical exponents close to 0.5 and a dimensionless tail close to 2.0, suggesting one-dimensional short-range structural disorder (Figure 6a) (52, 53). These estimates carried large uncertainties, driven mainly by the variance in ADC_K_ (TD), and should be regarded as suggestive. A Kärger exchange model fitted to K (TD) over 21 ≤ TD ≤ 250 ms and 37 ≤ TD ≤ 250 ms returned exchange times t_ex_ of 13.4 and 16.5 ms, consistent with the kurtosis-peak position (TD^∗^ = 21 ms) and with prior work (21, 54, 55); sampling more TDs below 21 ms would be needed to localize the kurtosis peak precisely (Figure 6b) (48).

The water DDES signal (Figure 6c), fitted with a tensor model, gave a *μ*FA of 0.53—well below the metabolite values (≈ 1.0)—indicating that water diffusion is multicompartmental and affected by exchange. Its mean diffusivity was lower than the diffusion-time ADC, consistent with diffusion-filtering effects at the higher DDES b-value, which potentially attenuates the faster, less-restricted (extracellular) signal pool.

## Discussion

The standard model of metabolite diffusion currently consists of randomly oriented cylinders (neurites) alone, and even this requires ultra-high b-values (> 10 ms µm^−2^) to be fitted consistently (5). Here we demonstrate a complementary strategy: multiparametric dMRS in the human brain, combining diffusion-time and double-diffusion encoding. Our data show a clear dependence of metabolite ADC and kurtosis on diffusion time, show that DDES provides complementary angular information, and demonstrate that jointly modeling both experiments extends the standard model with a second, soma-like compartment while removing the unstable, non-physical solutions that arise from either experiment alone.

### Metabolite diffusion and cellular morphometry

The diffusion-time dependence is consistent with restricting barriers—cell membranes and organelles—that increasingly hinder motion at longer time scales, and our ADC (TD) and K (TD) agree well with prior studies (Figure 3). At short TD (< 25 ms), metabolites probe the intracellular environment over short distances, giving higher ADC and lower kurtosis and reflecting cytoplasmic viscosity and molecular crowding. As TD increases, metabolites interact more with cellular boundaries, reducing the ADC toward a plateau three to four times below the intrinsic diffusivity D_0_ and raising the kurtosis. For neuronal markers (tNAA, Glu), the long-TD plateau was slightly below values reported in nonhuman primates, whereas for glial markers (tCho, mI) it matched primate values and exceeded rodent values. The latter is consistent with the greater morphological complexity of human and primate glia relative to rodents (18, 19, 56). The increase in K (TD) further supports increasing restriction and the non-Gaussian nature of metabolite diffusion in heterogeneous intracellular environments.

### Breaking the degeneracy with orthogonal encoding

Our central methodological finding is that the two-compartment inverse problem is degenerate when posed from diffusion-time data alone. The Monte-Carlo analysis exposed two near-indistinguishable solution branches—one non-physical (R_c_ > R_s_) and one plausible (R_c_ < R_s_)—with almost identical fitting curves over the measured TD range and differing chiefly in the back-extrapolated K (TD) at short, unmeasured diffusion times. Two routes can in principle resolve this ambiguity. The first is to measure K (TD) directly at short TD, which our experiments could not achieve because of PNS limits; this remains a promising option on ultra-strong-gradient systems. The second, which we pursued, is to add an orthogonal encoding. DDES alone could not provide accurate microstructure—its radial diffusivities and hence neurite radii tended to zero, reflecting coarse-graining at the comparatively long DDES diffusion time (27.3 ms)—but its intrinsic diffusivities reproduced the diffusion-time estimates on the R_c_ < R_s_ branch, suggesting that the two experiments carry consistent and complementary information.

Combining them bears this out. Initialized from four widely separated starting points (including the non-physical R_c_ > R_s_ branch), the joint fit converged stably to a single R_c_ < R_s_ solution, and the back-extrapolated K (TD) matched the constrained diffusion-time fit. The supporting evidence that metabolites are predominantly intra-neurite is internally consistent: *μ*FA ≈ 1 and MD ≈ D_0_ /3 are both expected signatures of stick-like diffusion. Excluding Glu, soma radii were slightly smaller in neurons (6.8 µm) than in glia (9.0 µm), with neurite radii of ≈1.3 µm in both. The equal neurite fractions (f_c_ of 0.82–0.90) across neuronal and glial markers are characteristic of primates and consistent with prior reports (18, 19).

### Relation to SANDI and the meaning of the estimated radii

Conceptually, our two-compartment description is the metabolite analogue of the soma-and-neurite density imaging (SANDI) model introduced for water diffusion (20), and recently translated to clinical hardware in combination with standard-model imaging (57). The crucial difference is specificity: water-based SANDI reports an apparent, compartment-averaged soma and neurite signal that mixes neuronal, glial, and—in gray matter—extracellular contributions, whereas metabolite dMRS isolates intracellular pools that are differentially enriched in neurons (NAA, Glu) and glia (tCho, mI). Multiparametric dMRS therefore offers cell-type–resolved soma and neurite estimates that water dMRI cannot, at the cost of lower SNR, single point resolution, and longer scans. The estimated radii should be interpreted as effective, model-dependent quantities rather than literal anatomical dimensions. Diffusion measurements have a well-characterized lower bound on the smallest resolvable radius, set by the available diffusion weighting, the noise floor, and orientation dispersion (58); moreover, the radius that diffusion MRI reports is an effective, tail-weighted quantity that overestimates the underlying mean radius, so small radii are systematically biased upward (59). This is relevant to our neurite radii (R_c_ ≈ 1–2.5 µm), which exceed the sub-micron radii of typical axons and dendrites and are best read as upper-bound, effective values; it is also consistent with the near-zero R_c_ recovered from DDES, where the long diffusion time removes sensitivity to small radii. For somata, the consistency of R_s_ across most metabolites (≈6–10 µm) is encouraging and physically reasonable, but the large values for Glu (and, in the diffusion-time–only fit, tCho) warrant caution.

In particular, the persistent Glu soma radius (≈20 µm) is unlikely to be biological. We considered whether non-neuronal glutamate pools could bias the estimate, but astrocytic glutamate constitutes only ≈ 10% and extracellular only 0.1% of the total tissue pool (60, 61). These fractions are too small to account for the anomalous Glu estimate. The most likely explanation is methodological as Glutamate is among the most challenging metabolites to quantify, owing to its strongly J-coupled multiplet, extensive spectral overlap (with glutamine, NAA, and macromolecules), and low effective SNR—particularly at the high b-values and long diffusion times. Our own data corroborate this: Glu showed a kurtosis decrease at the longest TD, the largest CRLBs in the DDES experiment, and two of four DDES datasets that had to be excluded. Therefore Glu microstructural estimates need to be treated with particular caution.

More broadly, we regard the neuron-versus-glia soma-size difference as a trend, not a statistically tested result, given the small DDES cohort and the Glu ambiguity.

### Water exchange and structural disorder

Metabolite cycling allowed water and metabolites to be measured in the same voxel and session, so the simultaneously acquired water signal provides a complementary, intracompartment-spanning view. Two features stand out. First, water K (TD) was non-monotonic, peaking near 21 ms—unlike the monotonic increase seen for metabolites. The Kärger model returned exchange times of 13–17 ms, in line with recent water-exchange estimates in gray matter (21, 54, 55). The contrast with the monotonic metabolite K (TD) is itself informative: it indicates that intercompartmental exchange is fast for water but negligible for metabolites on these time scales, which retrospectively justifies the non-exchanging assumption of our metabolite tissue model.

Second, the kurtosis peak position is sometimes used to estimate a characteristic restriction size; under the simplifying assumption that exchange is slow relative to coarse-graining, TD^∗^ ≈ 21 ms corresponds to a length scale of ≈10 µm. We caution, however, that this assumption is in tension with the short exchange time we estimate from the same data: the peak position is set jointly by restriction size and membrane permeability (10), so the ≈10 µm figure should be read as an order-of-magnitude estimate only.

The power-law analysis suggested one-dimensional short-range disorder, consistent with diffusion along neurites, but the large uncertainties mean this should be treated as qualitative. The lower water *μ*FA (0.53) relative to metabolites confirms that water samples multiple compartments with exchange, whereas metabolites remain confined to anisotropic intracellular spaces.

### Limitations

Several limitations should be emphasized.

#### (i) Two-cohort, cross-scanner design

The diffusion-time and DDES experiments were acquired in different cohorts, on different scanners, and at different ages (mean 33 vs. 57 years; mixed-sex vs. all-male). The joint fit therefore combines cohort-averaged curves from two populations rather than matched, within-subject data, and Figure 5 should be read as a proof of concept that orthogonal encoding breaks the model degeneracy, not as a definitive morphometric measurement. The principal evidence that the two datasets are comparable is the concordance of their intrinsic diffusivities D_0_ (Tables 1 and 2); nonetheless, a within-subject acquisition is an essential next step, and the cohort difference may contribute to some between-experiment discrepancies (e.g., in R_s_).

#### (ii) Uncertainty reporting

The joint-fit uncertainties (Figure 5) reflect convergence across starting conditions, not data variability; data-driven (bootstrap) uncertainties should accompany these estimates in future work.

#### (iii) T_2_-weighting and compartment filtering

Although prior work reports little effect of compartment-specific T_2_ over the echo-time ranges used here (5, 9, 21, 62, 63), we cannot fully exclude filtering effects whereby short-T_2_ restricted compartments are attenuated at the longer TE of semiLASER.

#### (iv) Residual orientational coherence

Despite maximizing gray-matter fraction in the PCC, a residual preferential fiber orientation cannot be excluded; this could introduce mild orientational anisotropy, particularly for the diffusion-time experiments, where gradient limits on the Connectom scanner precluded isotropic encoding. Future diffusion-time experiments should implement multidirectional encoding.

#### (v) Exchange-time sensitivity

The water exchange time was sensitive to data quality (e.g., the outlier-detection step), and additional TDs between 21 and 250 ms are needed to estimate it robustly.

#### (vi) Compartment assignment and glutamate reliability

The neuronal/glial assignment of metabolites is a useful approximation rather than a strict dichotomy. For glutamate specifically, however, the predominantly neuronal localization is well established (astrocytic 10%, extracellular 0.1%), so the anomalous Glu estimates are more plausibly attributed to its lower spectral reliability—J-coupling, spectral overlap, and low SNR at high b-value and long TD—than to compartment mixing.

### Clinical translation and outlook

The DDES arm was acquired entirely on clinical 3 T PRISMA hardware, and we have previously shown that short diffusion times (≈ 6 ms) are accessible on clinical scanners (14). While the full protocol used here also relied on the Connectom scanner for short-TD OG encoding and high-b STEAM kurtosis, the degeneracy-breaking benefit of combining diffusion-time and angular encoding does not require ultra-strong gradients. On ultra-strong-gradient systems, directly sampling K (TD) or the DDES signal at short diffusion times (< 10 ms) may resolve the model ambiguity without a separate angular experiment. Extending diffusion times beyond 500 ms could additionally constrain neurite length (12, 19, 42). For routine use, the protocol must be shortened by selecting a minimal set of diffusion times and rotation angles that preserves microstructural sensitivity and SNR; a candidate protocol of three diffusion times (6, 25, and 100 ms) and four DDES rotation angles (0, 90, 180, and 270°) should be feasible in well under one hour.

## Conclusion

Morphological changes in neurons and glia precede or accompany many neurological pathologies (64–67), motivating noninvasive techniques that can characterize cellular microstructure with cell-type specificity. We have presented the first multiparametric dMRS in humans, combining diffusion-time and double-diffusion encoding. The key methodological advance is that combining these orthogonal encodings breaks a fundamental degeneracy of single-encoding dMRS, allowing the standard neurite-only model to be extended with a soma compartment and stabilizing cellular modeling. Because dMRS isolates cell-type–enriched metabolites, the approach provides cell-type–specific soma and neurite estimates that water diffusion MRI cannot. Microglia and astrocytes change shape and size dramatically in response to immune challenges—spanning ramified (resting), hypertrophic (reactive), and amoeboid (phagocytic) states—and disentangling soma size from process complexity could improve disease phenotyping, for example in Alzheimer’s disease (67, 68). Although a clinically practical protocol still requires shortening and within-subject validation, multiparametric dMRS offers a concrete route toward quantitative, cell-type–specific microstructural imaging of the human brain.

## Author Contributions

A.D.: Conceptualization, Methodology, Software, Investigation, Formal analysis, Visualization, Writing—original draft. F.R.: Formal analysis, Methodology. K.Ş.: Software, Methodology (macromolecular baseline). M.A.: Methodology, Writing—review & editing. R.K.: Methodology (spectral fitting), Writing—review & editing. K.L.: Software (basis-set simulation), Writing—review & editing. D.K.J.: Resources, Funding acquisition, Writing—review & editing. J.V.: Methodology, Writing—review & editing. M.P.: Conceptualization, Methodology, Funding acquisition, Writing— review & editing.

## Data and Code Availability

All the scripts necessary to perform analysis are shared on (https://github.com/o-aNerd-o/Paper_2026_multiParaDiffMRS.git).

## Acknowledgements

We thank all the participants for contributing to this study and acknowledge the CIBM Center for Biomedical Imaging and sitem-insel Swiss Institute for Translational and Entrepreneurial Medicine for providing expertise and resources.

