## Supplementary Materials for "Resolving Cellular Morphology in the Human Brain with Multiparametric Diffusion MR Spectroscopy"

### Supporting Information for: Resolving Cellular Morphology in the Human Brain with Multiparametric Diffusion MR Spectroscopy

#### Experimental setup

Experiments were conducted at two sites on three Siemens MR Scanners (Connectom-A, 2×PRISMA). Table S1 provides an overview of the scanner hardware and software configurations, and Table S2 details the sequences used.

Table S1: MR Scanner hardware and software

| | | |
|---|---|---|
| **Cardiff University Brain Research Imaging Centre (CUBRIC, Cardiff)** | | |
| a. | Field strength | 3 T |
| b. | Manufacturer | Siemens |
| c. | Model | Connectom-A, VD11D |
| d. | RF coils | $^1$H, 32 receive channel, head |
| **Cardiff University Brain Research Imaging Centre (CUBRIC, Cardiff)** | | |
| a. | Field strength | 3 T |
| b. | Manufacturer | Siemens |
| c. | Model | Prisma, VE11C |
| d. | RF coils | $^1$H, 32 receive channel, head |
| **Swiss Institute for Translational and Entrepreneurial Medicine (SITEM, Bern)** | | |
| a. | Field strength | 3 T |
| b. | Manufacturer | Siemens |
| c. | Model | Prisma, VE11C |
| d. | RF coils | $^1$H, 32 receive channel, head |

More details on the experimental conditions (number of averages and sample sice, rotation angles in DDES) are provided in the main manuscript.

Table S2: dMRS Sequences

| | | | | |
|---|---|---|---|---|
| **Diffusion-time experiments** (TD = 6 . . . 50ms), **Connectom-A** | | | | |
| a. | Pulse sequence | semiLASER | | |
| b. | TE/TR | 76/3000 ms (2 prep. scans) | | |
| c. | Diffusion settings | Parameters | Diffusion Time | $b_{max}$ |
| | OG | sine modulated ($2 \times 20.5$ ms, $\alpha$/N: 1/0) | 6.0 ms | $3\,\mathrm{ms\,\mu m^{-2}}$ |
| | PG | $\Delta/\delta/\epsilon$: 21/7.1/3.2 ms | 18.6 ms | $3\,\mathrm{ms\,\mu m^{-2}}$ |
| | PG | $\Delta/\delta/\epsilon$: 37/7.1/3.2 ms | 34.6 ms | $3\,\mathrm{ms\,\mu m^{-2}}$ |
| **Diffusion-time experiments** (TD = 50 . . . 250ms), **Connectom-A** | | | | |
| a. | Pulse sequence | STEAM, 6 OVS bands on each side | | |
| b. | TE/TM/TR | 37/32;82;232/3000 ms (2 prep. scans) | | |
| c. | Diffusion settings | Parameters | Diffusion Time | $b_{max}$ |
| | PG | $\Delta/\delta/\epsilon$: 50/7.1/3.2 ms | 47.6 ms | $8\,\mathrm{ms\,\mu m^{-2}}$ |
| | PG | $\Delta/\delta/\epsilon$: 100/7.1/3.2 ms | 97.6 ms | $8\,\mathrm{ms\,\mu m^{-2}}$ |
| | PG | $\Delta/\delta/\epsilon$: 250/7.1/3.2 ms | 247.6 ms | $8\,\mathrm{ms\,\mu m^{-2}}$ |
| both experiments were performed in the same session | | | | |
| d. | VOI | Midline PCC (GM/WM/CSF $70.9 \pm 1.7/16.5 \pm 2.6/12.7 \pm 3.3\%$) | | |
| e. | Nominal VOI size | $10.0 \pm 1.5$mL | | |
| **Double-diffusion encoding experiments, PRISMA** | | | | |
| a. | Pulse sequence | semiLASER, 2 OVS bands along excitation direction | | |
| b. | TE/TR | 125/2500 ms (4 prep. scans) | | |
| c. | Diffusion settings | Parameters | Diffusion Time | $b_{max}$ |
| | DDE | $\Delta/\delta/\epsilon/\tau$: 32/14/1/39 ms | 27.3 ms | $5.2\,\mathrm{ms\,\mu m^{-2}}$ |
| d. | VOI | Midline PCC (GM/WM/CSF $63.0 \pm 5.4/22.7 \pm 5.0/14.3 \pm 0.8\%$) | | |
| e. | Nominal VOI size | $16.1 \pm 2.4$mL | | |
| f. | Water suppression | no WS; metabolite Cycling | | |

### Sequence validation

The results from sequence validation are presented in Fig. S1. Irrespective of sequence type or diffusion encoding strategy, phantom measurements yield equal diffusivities for metabolites (B, diffusion-time experiment; D, double-diffusion encoding experiment) and water (A, diffusion-time experiment). More detailed information is provided in the main manuscript.

Moreover, we investigated potential effects from spurious echos in STEAM. Fig. S1A shows the signal amplitudes in the NIST phantom for the different sequence configurations, with equal amplitudes for semiLASER (equal TEs), but lower for STEAM, even though TE is shortened by a factor of two. T1 and T2 are predefined by the NIST in the vial with 40 % PVP at values of 710 ms and 550 ms, respectively ($D = 0.56\,\mathrm{\mu m^2\,ms^{-1}}$). The T1 and T2 corrected signal gain $g$ of spin (SE) over stimulated (STE) echo is given by:

$$\frac{S^{SE}}{S^{STE}} = g \cdot \frac{e^{-\frac{TE^{SE} - TE^{STE}}{T_2}}}{e^{-\frac{TM^{STE}}{T_1}}} \tag{1}$$

where $\mathrm{TE^{SE}}$ and $\mathrm{TE^{STE}}$ denote the echo times for semiLASER and STEAM, and $\mathrm{TM^{STE}}$ the mixing time for STEAM. Relying on the water signal amplitudes at the lowest b-value in Fig. S1A, this yields a signal gain factor of $g = 1.83$ of semiLASER over STEAM, which is close to the theoretical value of $g = 2$. For sequence validation experiments no OVS bands were set initially. Indeed, repeating the STEAM measurement at the lowest b-value with 6 OVS bands enclosing the voxel on each side $g = 1.96$ was found. The higher g factor most likely originates from suppressed out-volume signals in STEAM with OVS applied and, in turn, we used six OVS bands for the *in vivo* experiments.

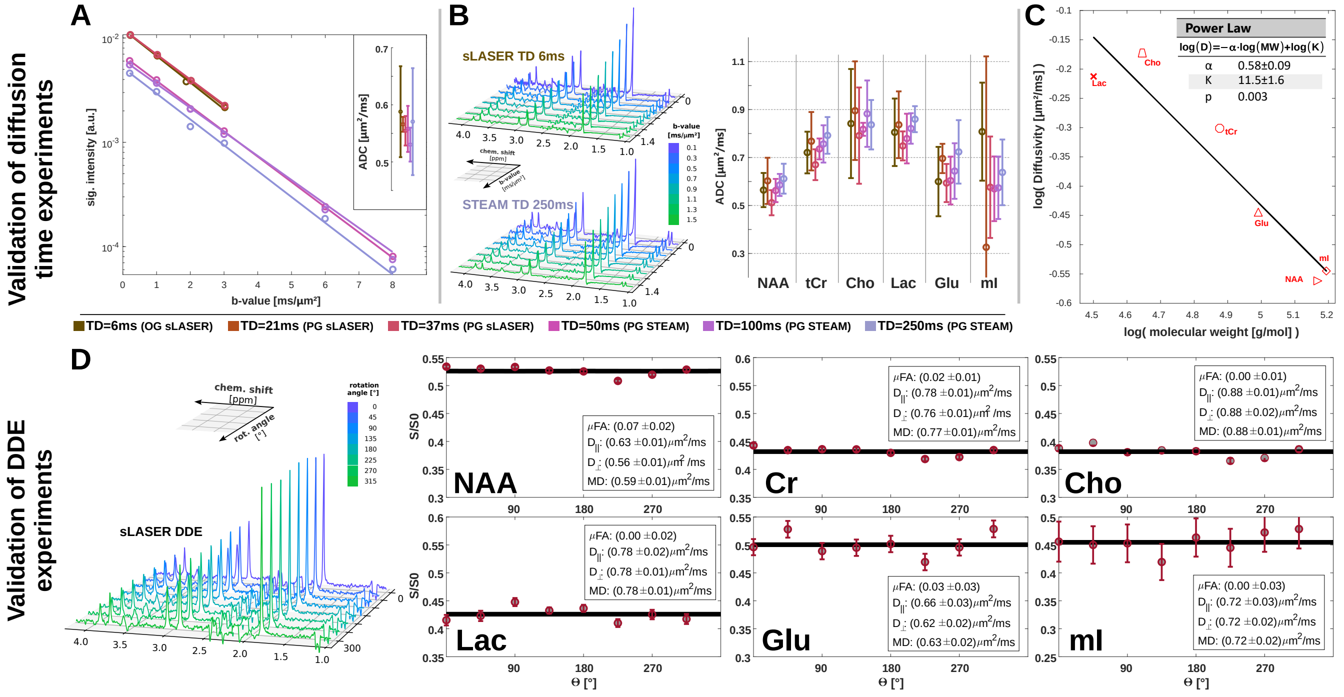


Figure S1: **Sequence validation**: Diffusion-time experiments were validated in a NIST (A) and Braino (B) phantom for the applied sequences and sequence settings (echo-type (STEAM vs semiLASER), echo-time (37 vs 76ms) and diffusion-encoding (pulsed- vs oscillating-gradients)). Validation of the double-diffusion-encoding (D) used free metabolite diffusion the Braino phantom. The estimated diffusivities are equal, irrespective of diffusion encoding direction ($D_{\parallel} = D_{\perp} = MD$) and agree with results from the diffusion-time experiments. The estimated free diffusivities follow a power-law (C) with respect to the molecular weights of the metabolites.

### Spectral Quality Measures

23

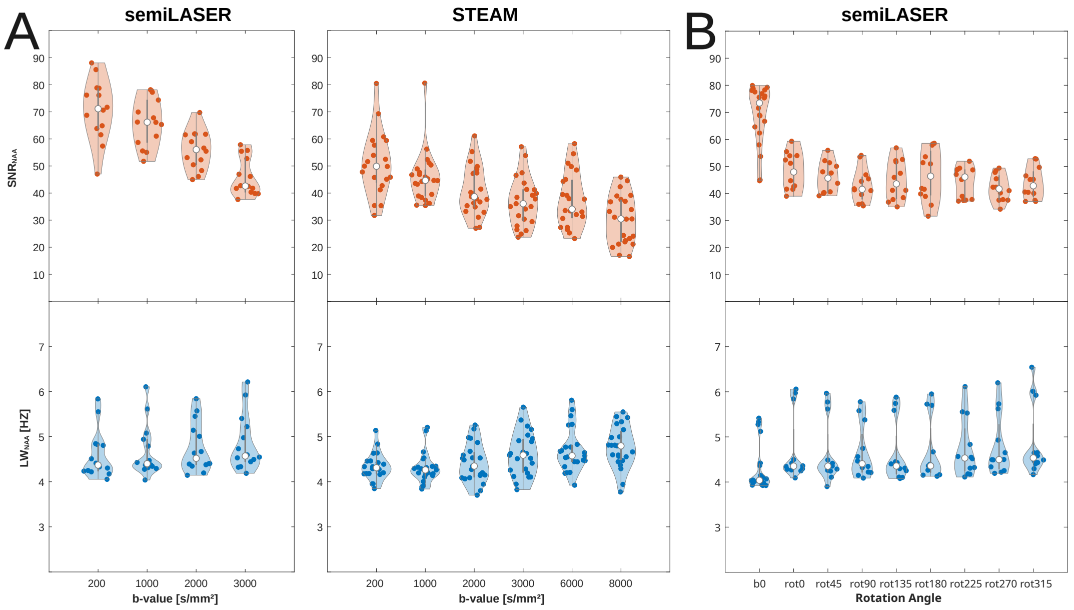


Figure S2: **Quality Measures**: Signal-to-noise-ratio (SNR) and linewidth (LW) obtained in the *in vivo* experiments from fitting the NAA singlet for the diffusion-time (A) and double-diffusion encoding (B) experiments combined over all subjects, diffusion-times, individual b-values, and rotation angles.

### Monte Carlo Simulation of Diffusion Time dependence Modelling

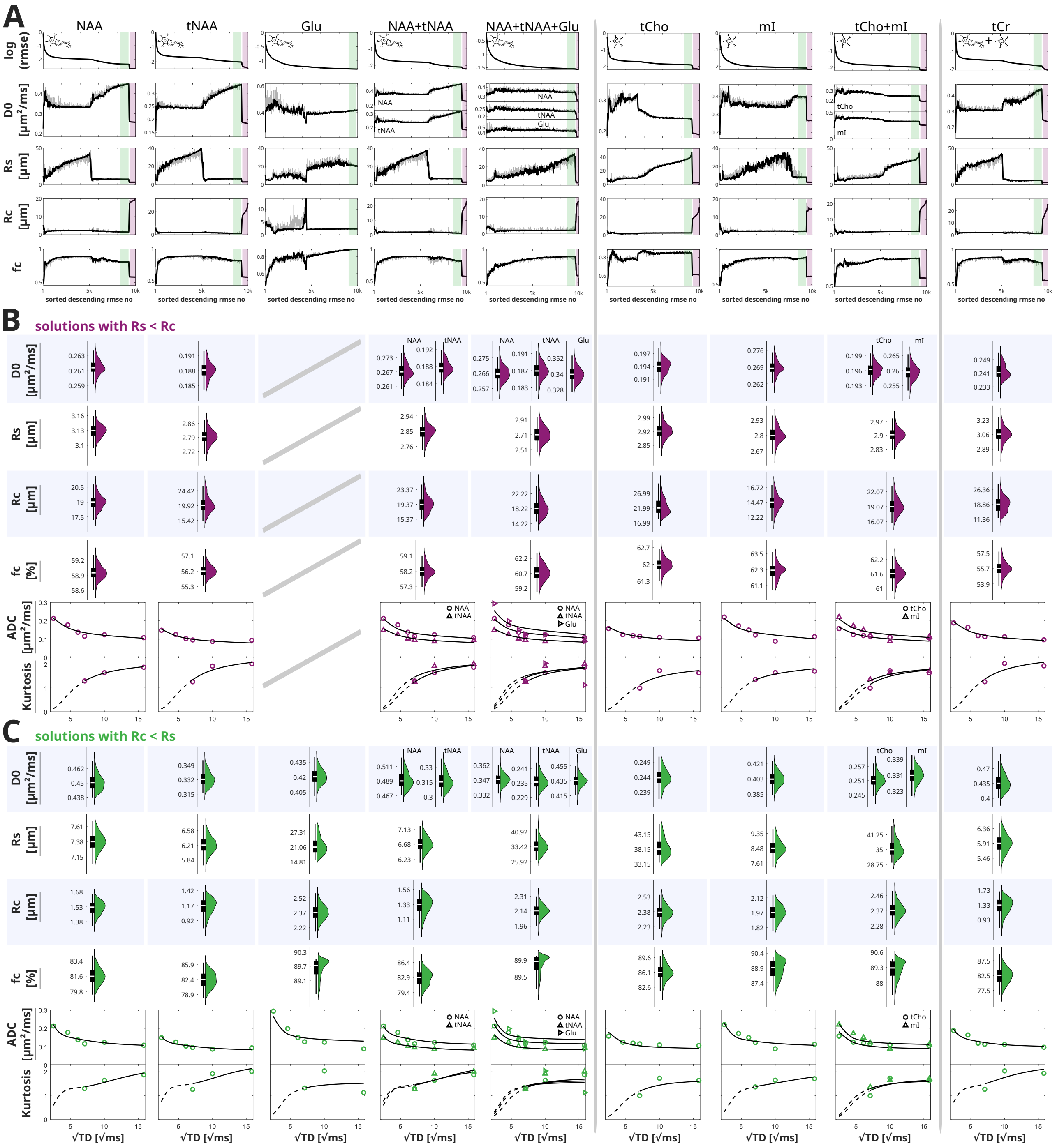


Figure S3: **Monte Carlo simulation of the diffusion-time dependence implying a two-compartment model:** (A) shows the result from fitting 10000 randomly initialized starting parameters for the intrinsic diffusivity $D_0$, sphere radius $R_s$ (soma radius), cylinder radius $R_c$ (radius of neurites), and cylinder fraction $f_c$ sorted descending by the lowest root-mean-square-error (RMSE) for neuronal markers (NAA, tNAA, Glu), glial markers (tCho, mI), tCr and their joint fitting (joint fitting considers metabolites inhabiting the same compartment where $R_s$, $R_c$ and $f_c$ were set as global parameters, and metabolites intrinsic diffusivities $D_0$ were fitted separately). The green areas indicate the 10% quantile of the lowest RMSEs with $R_c < R_s$, and the red areas the 90% quantile of the lowest RMSEs with $R_s < R_c$. (B) and (C) shows the histograms of the estimated microstructural properties within these quantile limits for the first plateau in green and the second in red. Bellow, the corresponding fitting results of ADC(TD) and K(TD) using the median values within the respective quantile limits (dashed lines indicate the back-estimated K(TD)).

Fig. S3 presents the stability analysis of the modelling of the diffusion-time dependence. Monte Carlo simulations used 10 000 randomly initialized starting parameters. The root-mean-square-errors (RMSEs) indicate an ambiguous solution space responsible for an unstable fitting.

We found that RMSEs extend over a wide range of several orders of magnitude, indicating that the fitting routine is not able to approach a stable minimum. Although overall RMSE values converge, reaching a first plateau with $R_c < R_s$ and subsequently a second well-separated plateau with $R_s > R_c$ (except for Glu where $R_c$ remained smaller than $R_s$), estimated model parameters on each plateau show high levels of variance. This behaviour is found consistently for all metabolites. Moreover, the solutions at the second plateau, even though at lower RMSEs, contradict realistic cellular morphologies with larger soma than neurite radii. The estimated free diffusivities ($D_0$) and cylinder fractions ($f_c$) for these solutions ($R_c > R_s$, red in Fig. S3B) are by average 68.5 % and 69.6 % lower than those on the first plateau ($R_c < R_s$, green in Fig. S3C). Also, note the difference in the diffusion-time course of K(TD) back-estimated towards shorter diffusion times (dashed lines) between (B) and (C). The detailed results from refitting the diffusion-time data with starting conditions constrained to $R_c < R_s$ are presented in Fig. S4.

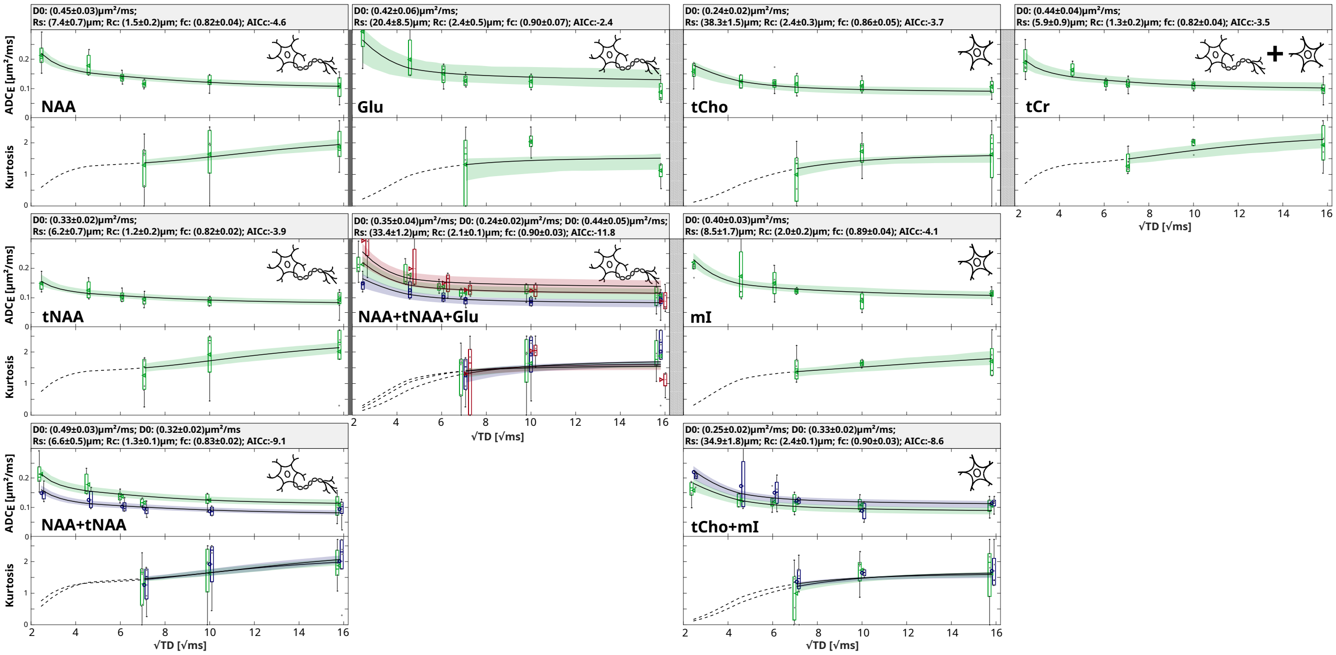


Figure S4: **Refit of diffusion-time dependence:** Fitting results of the diffusion-time dependence ADC(TD) and K(TD) using the median of the solutions from Monte Carlo simulations with $R_c < R_s$ as starting values.

### Multiparametric diffusion modeling

Multiparametric diffusion modeling combined diffusion-time and double-diffusion encoding and was tested by using four initial sets of starting conditions from the MC simulations: (i) the median values from the biophysical solutions with $R_c < R_s$ (green in Fig. S3, equal to the starting conditions used for refitting of diffusion-time dependence), (ii) the median values from the solutions that contradict realistic cellular morphology with $R_c > R_s$ (lowest RMSEs, red in Fig. S3), (iii) the solution at the highest RMSEs with index #1, and (iv) a solution where $D_0$, $R_s$, and $R_c$ show a strong gradient towards the next lower RMSE value. The results from fitting are presented in Tab. S3.

Table S3: **Multiparametric Modelling:** Summary of the four initial test conditions and fitting results using joint modelling of diffusion-time and double-diffusion encoding.

| | | NAA | | | | tNAA | | | | Glu | | | |
|---|---|---|---|---|---|---|---|---|---|---|---|---|---|
| | Condition | (i) Rc<Rs | (ii) Rc>Rs | (iii) | (iv) | (i) Rc<Rs | (ii) Rc>Rs | (iii) | (iv) | (i) Rc<Rs | (ii) Rc>Rs | (iii) | (iv) |
| Starting Con. | D0 [µm²/ms] | 0.45 | 0.26 | 0.23 | 0.46 | 0.33 | 0.19 | 0.15 | 0.24 | 0.42 | 0.38 | 0.25 | 0.44 |
| | Rs [µm] | 7.29 | 3.13 | 11.27 | 21.74 | 6.17 | 2.79 | 11.64 | 38.53 | 21.77 | 7.86 | 10.14 | 9.32 |
| | Rc [µm] | 1.59 | 19.04 | 5.28 | 0.52 | 1.27 | 20.45 | 6.39 | 1.91 | 2.37 | 8.93 | 8.92 | 2.11 |
| | fc | 0.82 | 0.59 | 0.55 | 0.8 | 0.82 | 0.56 | 0.6 | 0.89 | 0.89 | 0.74 | 0.64 | 0.79 |
| Fitting Result | D0 [µm²/ms] | 0.47 | 0.47 | 0.47 | 0.48 | 0.33 | 0.33 | 0.34 | 0.35 | 0.45 | 0.45 | 0.45 | 0.46 |
| | Rs [µm] | 7.35 | 7.35 | 7.35 | 7.27 | 6.37 | 6.21 | 6.19 | 6.28 | 20.25 | 20.25 | 21.75 | 20.09 |
| | Rc [µm] | 1.44 | 1.44 | 1.44 | 1.40 | 1.06 | 1.17 | 1.07 | 0.72 | 2.16 | 2.16 | 2.19 | 2.11 |
| | fc | 0.83 | 0.83 | 0.83 | 0.83 | 0.80 | 0.82 | 0.81 | 0.79 | 0.90 | 0.90 | 0.90 | 0.90 |

| | | tCho | | | | mI | | | | tCr | | | |
|---|---|---|---|---|---|---|---|---|---|---|---|---|---|
| | Condition | (i) Rc<Rs | (ii) Rc>Rs | (iii) | (iv) | (i) Rc<Rs | (ii) Rc>Rs | (iii) | (iv) | (i) Rc<Rs | (ii) Rc>Rs | (iii) | (iv) |
| Starting Con. | D0 [µm²/ms] | 0.24 | 0.19 | 0.18 | 0.32 | 0.4 | 0.27 | 0.18 | 0.46 | 0.43 | 0.24 | 0.16 | 0.31 |
| | Rs [µm] | 38.73 | 2.92 | 9.78 | 10.13 | 9.04 | 2.8 | 13.34 | 7.42 | 5.7 | 3.05 | 15.21 | 39.22 |
| | Rc [µm] | 2.41 | 22.78 | 5.27 | 1.29 | 2.05 | 14.59 | 6.66 | 0.86 | 1.54 | 18.78 | 6.3 | 2.01 |
| | fc | 0.86 | 0.62 | 0.51 | 0.77 | 0.89 | 0.62 | 0.61 | 0.75 | 0.89 | 0.56 | 0.61 | 0.89 |
| Fitting Result | D0 [µm²/ms] | 0.36 | 0.36 | 0.34 | 0.35 | 0.45 | 0.45 | 0.45 | 0.47 | 0.44 | 0.44 | 0.45 | 0.42 |
| | Rs [µm] | 9.13 | 9.13 | 11.27 | 10.11 | 8.29 | 8.29 | 8.27 | 7.52 | 6.18 | 5.91 | 6.18 | 5.77 |
| | Rc [µm] | 1.01 | 1.01 | 1.15 | 1.16 | 1.80 | 1.80 | 1.81 | 1.81 | 1.05 | 1.33 | 1.10 | 1.50 |
| | fc | 0.81 | 0.81 | 0.82 | 0.82 | 0.90 | 0.90 | 0.90 | 0.90 | 0.76 | 0.82 | 0.79 | 0.83 |

### Data

Table S4: **Diffusion-Time Data:** Cohort results of estimated $ADC_E(TD)$ and $K(TD)$ values. Uncertainties are derived from the standard deviation over all subjects.

| | TD | NAA ADC$_E$ [µm²/ms] | NAA K | tNAA ADC$_E$ [µm²/ms] | tNAA K | Glu ADC$_E$ [µm²/ms] | Glu K |
|---|---|---|---|---|---|---|---|
| neuronal | 6 | 0.21±0.05 | / | 0.15±0.02 | / | 0.29±0.08 | / |
| | 21 | 0.18±0.05 | / | 0.13±0.04 | / | 0.20±0.09 | / |
| | 37 | 0.14±0.01 | / | 0.10±0.02 | / | 0.15±0.04 | / |
| | 50 | 0.12±0.01 | 1.29±0.78 | 0.10±0.02 | 1.26±0.61 | 0.13±0.02 | 1.31±1.07 |
| | 100 | 0.12±0.02 | 1.64±0.95 | 0.09±0.02 | 1.92±0.98 | 0.12±0.03 | 2.04±0.33 |
| | 250 | 0.11±0.04 | 1.87±0.59 | 0.09±0.03 | 2.01±0.91 | 0.09±0.04 | 1.12±0.38 |

| | TD | tCho ADC$_E$ [µm²/ms] | tCho K | mI ADC$_E$ [µm²/ms] | mI K | | tCr ADC$_E$ [µm²/ms] | tCr K |
|---|---|---|---|---|---|---|---|---|
| glial | 6 | 0.16±0.04 | / | 0.22±0.06 | / | unspecific | 0.19±0.05 | / |
| | 21 | 0.12±0.03 | / | 0.17±0.11 | / | | 0.16±0.03 | / |
| | 37 | 0.12±0.03 | / | 0.15±0.05 | / | | 0.12±0.02 | / |
| | 50 | 0.12±0.04 | 0.99±0.80 | 0.12±0.01 | 1.36±0.65 | | 0.11±0.02 | 1.26±0.56 |
| | 100 | 0.11±0.03 | 1.72±0.52 | 0.09±0.04 | 1.64±0.11 | | 0.11±0.02 | 2.04±0.29 |
| | 250 | 0.11±0.03 | 1.63±1.00 | 0.11±0.02 | 1.71±0.62 | | 0.10±0.03 | 1.93±0.65 |

Table S5: **Double-diffusion encoding Data:** Cohort results of estimated DDES(Θ) values, with signal at $b = 0$ normalized to 1. Uncertainties are derived from the standard deviation over all subjects (for mI one, for Glu two valid dataset, and four otherwise).

| | Θ | NAA DDE(Θ) | tNAA DDE(Θ) | Glu DDE(Θ) |
|---|---|---|---|---|
| neuronal | 0 | 0.58±0.01 | 0.62±0.01 | 0.52±0.03 |
| | 45 | 0.53±0.03 | 0.59±0.03 | 0.47±0.03 |
| | 90 | 0.50±0.02 | 0.56±0.02 | 0.43±0.01 |
| | 135 | 0.52±0.03 | 0.57±0.02 | 0.48±0.05 |
| | 180 | 0.56±0.02 | 0.60±0.02 | 0.51±0.01 |
| | 225 | 0.53±0.01 | 0.58±0.02 | 0.49±0.02 |
| | 270 | 0.50±0.01 | 0.56±0.02 | 0.48±0.10 |
| | 315 | 0.54±0.03 | 0.60±0.03 | 0.51±0.03 |

| | Θ | tCho DDE(Θ) | mI DDE(Θ) | | tCr DDE(Θ) |
|---|---|---|---|---|---|
| glial | 0 | 0.65±0.01 | 0.71 | unspecific | 0.60±0.01 |
| | 45 | 0.62±0.03 | 0.59 | | 0.56±0.03 |
| | 90 | 0.59±0.02 | 0.58 | | 0.54±0.02 |
| | 135 | 0.6±0.03 | 0.64 | | 0.55±0.03 |
| | 180 | 0.63±0.03 | 0.65 | | 0.58±0.03 |
| | 225 | 0.62±0.02 | 0.63 | | 0.56±0.02 |
| | 270 | 0.60±0.02 | 0.54 | | 0.54±0.02 |
| | 315 | 0.63±0.02 | 0.62 | | 0.57±0.03 |
